# A microfluidic gradient and parallel-track system uncovers spatial control of endocytosis and adhesion formation in breast cancer cell migration

**DOI:** 10.1101/2025.06.24.661139

**Authors:** Emily T. Chan, Travis H. Jones, Cristopher M. Thompson, Hariharan Kannan, Malcolm W. D’Souza, Mushtaq M. Ali, Cömert Kural, Jonathan W. Song

**Author notes:** Corresponding authors: Cömert Kural, Jonathan Song.

## Abstract

Cell migration through confined spaces is a critical step in cancer metastasis, yet the spatial regulation of endocytosis and adhesion dynamics during this process remains poorly understood. To address this, we developed a microfluidic platform that generates stable, spatially linear biochemical gradients across 5 μm-tall migration channels while limiting confounding flow-induced shear stress (<0.05 dyn/cm^2^). COMSOL simulations and optical calibration using FITC-dextran confirmed that gradients form reliably within 5 minutes. The microdevice also supports long-term live imaging and is compatible with both spinning disk confocal and total internal reflection fluorescence structured illumination microscopy modalities, enabling high-resolution visualization of adhesion and endocytic structures. Localized application of the endocytic inhibitor Dyngo-4a to the front or rear of migrating cells revealed that front-targeted inhibition significantly increased the enrichment of paxillin and the clathrin adaptor AP-2 at the leading edge, whereas rear-targeted inhibition completely abolished their front-rear asymmetry. These changes were accompanied by enhanced migration speed and persistence, particularly under front-targeted inhibition. Together, these findings highlight the critical role of spatially coordinated endocytosis in sustaining polarized adhesion and persistent cell movement. Our platform offers a powerful tool for dissecting subcellular mechanisms of migration under confinement and provides a broadly applicable framework for probing spatially localized signaling in engineered microenvironments.

## Introduction

Cell migration is central to numerous physiological processes including immune surveillance (*1*), wound healing (*2*), and embryonic development (*3*). In cancer, however, similar migratory machinery is co-opted to drive tumor invasion and metastasis (*2*, *4*), requiring cells to traverse narrow, confined spaces within the extracellular matrix (ECM) (*2*, *4*, *5*). These spatial constraints dramatically alter cell morphology, cytoskeletal architecture, and intracellular signaling (*5–7*). Despite the biological and clinical importance of confined cell migration, how cells integrate spatial biochemical cues with physical confinement to coordinate motility remains incompletely understood.

A key but underexplored component of this integration is the spatial regulation of membrane trafficking processes such as endocytosis. Clathrin-mediated endocytosis (CME) orchestrates key events during cell migration, including receptor internalization and focal adhesion (FA) turnover (*8–10*). Prior studies show that CME and FAs exhibit spatial asymmetries: CME is often enriched at the leading edge and suppressed ventrally, while mature FAs accumulate toward the rear of migrating cells (*11*, *12*). Yet, it remains unclear how these polarized distributions are maintained or altered under physical confinement. Most conventional systems lack the spatial resolution and environmental control to probe these dynamics at subcellular scales, especially in live, migrating cells.

Dissecting the region-specific roles of endocytosis in migration requires spatially targeted, rather than global, perturbations. Endocytosis is inherently polarized in migrating cells, with distinct roles at the front and rear (*13*). To enable such investigations, an experimental platform must 1) establish a stable, spatially linear chemotactic gradient along the cell axis, 2) support high-resolution imaging of subcellular structures, and 3) permit selective delivery of perturbations, such as pharmacological inhibitors, to defined regions of the cell. Few existing systems satisfy all these criteria simultaneously.

In this study, we introduce a microfluidic device engineered to meet these demands. Our platform generates reproducible chemotactic gradients within narrow, cell-sized channels that minimize shear stress and maintain chemical stability. It is compatible with both confocal and super-resolution microscopy, enabling live visualization of CME and FA dynamics with subcellular resolution. Crucially, the system allows spatially restricted delivery of a CME inhibitor to either the front or rear of migrating breast cancer cells, which has not been previously demonstrated in confined chemotaxis assays. Using this platform, we uncover that front- and rear-targeted CME inhibition elicits distinct effects on adhesion localization, cell polarity, and migratory behavior. Our findings underscore the importance of spatial membrane dynamics in confined migration and demonstrate how microscale engineering can resolve previously inaccessible questions in cancer metastasis.

## Methods

### Cell lines and reagents

Genome-edited AP2-eGFP expressing human breast cancer SUM159 cell lines (eGFP incorporated at the C-terminus of the σ2 subunit of AP-2, a marker for endocytic clathrin structures) were kindly gifted by Dr. Tomas Kirchhausen (Harvard Medical School). SUM159 cells were cultured in complete media consisting of phenol red-free Ham’s F-12 (Caisson Labs, HFL05), supplemented with 5% fetal bovine serum (Gibco, A5256701), 1% penicillin and streptomycin (Thermo Fisher Scientific Inc., 15140122), 1 μg/mL hydrocortisone (Sigma-Aldrich, H-4001), 5 μg/mL insulin (Cell Applications, 128-100), and 10 mM 4-(2-hydroxyethyl)-1-piperazine-ethane-sulfonic acid (HEPES, Corning Inc., 25 -060-CI) at pH 7.4. All cells were maintained at 37 °C and 5% CO_2_ and passaged every 2-3 days. Each cell stock solution was replaced after a maximum of 8 passages.

### Design and fabrication of microfluidic migration devices

The micron-scale and multi-level features of the microfluidic migration device were first patterned onto silicon wafers using SU-8 photoresist (*14*). The fabricated device dimensions are designed as follows: 30 migration channels (5 μm tall, 20 μm wide, 150 μm long); 2 chambers (100 μm tall, 100 μm wide, 1400 μm long); 2 inlets and 1 outlet (radii 375 μm). Fabrication of multi-level features required two transparency photomasks designed in AutoCAD 2021: one defining the thicker chamber, inlets, and outlet regions at approximately 100 μm in height, and the other defining the thinner migration channels at approximately 5 μm in height. To create the master mold, a silicon wafer was spin-coated with negative SU-8 photoresist, where SU-8 2100 (Kayaku Advanced Materials, Inc., Y111075 0500L1GL) was used to form the thick features and SU-8 2005 (Kayaku Advanced Materials, Inc., Y111045 0500L1GL) was used for the thin channels. The wafer was exposed to UV light using a maskless aligner (Heidelberg Instruments, MLA 150) to ensure precise alignment of the two layers. After UV exposure, the SU-8 2100 layer was developed using SU-8 developer (Kayaku Advanced Materials, Inc., Y020100 4000L1PE), generating the negative relief of the design. The wafer underwent a hard bake at 150 °C for 2 hours to eliminate cracks and refine structural details. To prevent polydimethylsiloxane (PDMS) from adhering to the mold, silanization was performed by passivating the wafer with tridecafluoro-1,1,2,2-tetrahydrooctyl-1-trichlorosilane (United Chemicals Ltd, T2492) for 30 minutes in a fume hood.

The developed SU-8 master was then placed into a 150 mm petri dish (VWR, 10062-882) for PDMS replica molding (*15*). A 10:1 mixture of PDMS base elastomer to cross-linker (Dow Corning Corporation, Sylgard 184 Silicone Elastomer) was poured over the mold, degassed, and cured at 65 °C for 2 hours. The cured PDMS layer, approximately 0.55-0.65 cm thick, was peeled off and access ports were introduced by punching a 0.5 mm hole at the outlet using a stainless steel biopsy punch (Robbins Instruments, RBP-05) and two 4 mm inlet holes using a biopsy punch (Integra LifeSciences Services, 33-34-P/25). These inlets were positioned symmetrically to ensure proper fluid distribution. The PDMS layer was irreversibly bonded to a No. 1.5 glass coverslip inside a 35 mm dish (MatTek, P35G-1.5-20-C) using plasma oxidation (30W, 650 mbar, 70 seconds; Harrick Plasma, PDC-001) followed by baking at 65 °C for 10 minutes. This multi-layer assembly process finalized the formation of the microfluidic migration chamber.

### Validation of dimensions of migration devices

Microstructural features of the SU-8 microfluidic migration device were characterized using scanning electron microscopy (SEM) at Ohio State University’s Nanotech West Laboratory. A row of devices patterned on the wafer was fractured at specific positions to expose channel sidewalls for cross-sectional imaging. Prior to mounting, a thin layer of gold-palladium was deposited on the broken wafer pieces using a Cressington 108 Auto/SE Sputter Coater to reduce surface charging and enhance image contrast.

After sputter coating, samples were affixed to aluminum pin stubs using conductive carbon tape, and electrical grounding was further ensured using colloidal silver paste. SEM imaging was conducted using a Zeiss Ultra Plus field emission SEM. Samples were introduced via the loadlock chamber and mounted on a single stub mount holder. Images were acquired under high vacuum using the in-lens secondary electron detector at an accelerating voltage of 5.0 kV, a stage tilt of 45°, and a working distance of approximately 4.2-2.5 mm. Imaging was performed at 250-5000X magnification with a 60 μm aperture and a frame acquisition time of 15.9 seconds to capture the full cross-section of microchannels and assess their height, width, and sidewall integrity.

### Characterization of migration devices

Fluid dynamics and molecular transport within the migration device were modeled using COMSOL Multiphysics 6.2 (COMSOL Inc.), employing the CFD and Transport of Diluted Species modules. The device geometry was reconstructed based on experimental dimensions: migration channels measuring 5 μm (H) × 20 μm (W) × 150 μm (L), two chambers measuring 100 μm (H) × 100 μm (W) × 1400 μm (L), and inlets and outlets with radii of 375 μm and 250 μm, respectively. Default properties of water at 37°C were used to approximate phosphate-buffered saline (PBS).

A steady-state creeping flow model was implemented to simulate low-shear conditions relevant to cell migration. Boundary conditions included no-slip conditions along the walls, no-flux boundaries except at the two inlets and single outlet, and an outlet set to atmospheric pressure. A constant withdrawal rate of 5 µL/h was applied at the outlet, ensuring equal flow input from both inlets. To simulate molecular transport, a time-dependent diffusion model was solved for 10 kDa FITC-dextran, using a diffusion coefficient of 1.43 × 10^-10^ m^2^/s calculated via the Stokes-Einstein equation (*16*). The initial concentration was set to 0.1 mol/m^3^ in the source chamber and 0 mol/m^3^ in the sink chamber. Simulations were run for 5 hours to analyze gradient formation and stability across the migration channels.

Fluorescence intensity profiles along the migration channels were extracted and normalized to assess gradient formation over time. The final steady-state concentration distributions were compared across channels at multiple, defined time points. Mesh refinement was performed to ensure numerical stability and minimize computational error.

### Gradient characterization in migration devices

A chemotactic gradient in the migration device was characterized using 10 kDa FITC-dextran (Sigma-Aldrich, FD10S) and imaged with a spinning disk confocal microscope. After fabrication and bonding of the device to a No. 1.5 glass coverslip (Fisherbrand, 12544E), a stainless steel pin (New England Small Tube, NE-1310-03) was bent and connected to plastic tubing (0.020” ID x 0.060” OD Tygon® ND 100-80 Microbore Tubing, 56515), approximately 0.5 meters in length. The tubing was threaded through a drilled 0.5 cm diameter hole in a 35 mm plastic dish lid (MatTek, P35G-1.5-20-C) and connected to a 2.5 mL glass syringe (Hamilton Company, 81420) fitted with a metal needle hub (Hamilton Company, 7748-09). The syringe was mounted on a syringe pump (Harvard Apparatus, 70-3007) to facilitate media infusion and withdrawal. Before attachment to the microdevice, the syringe, needle, tubing, and stainless steel pin were pre-filled with 1X PBS (Corning, 21-031-CV). The pin was connected to the device outlet, and the syringe pump was set to infuse at 5 μL/h to establish stable flow within the device.

After securing the device on the confocal microscope stage, 50-60 μL of 1X PBS (Thermo Fisher Scientific Inc., 10010023) was added to each inlet. To equilibrate pressures between the two inlets, the syringe pump was switched to withdrawal mode at 5 μL/h for 1.5 hours. Two pipettes were then used to simultaneously introduce equal volumes of 1X PBS and a concentrated solution of FITC-dextran into the respective inlets (less than 2 μL each). The FITC-dextran solution was prepared at a concentration that would yield a final inlet concentration of 1 mg/mL, accounting for an estimated 3.75 μL reduction per inlet volume due to prior flow. Following FITC-dextran introduction, a continuous withdrawal rate of 5 μL/h was maintained for up to 5 hours during imaging.

FITC-dextran fluorescence was captured using a spinning disk confocal microscope (see **Spinning disk confocal image acquisition**) with 488 nm excitation (300 ms exposure time, 25% power) every 1 minute for 7 hours using a 20X objective lens (NA 0.75, Nikon) at 37 °C. The Nikon Perfect Focus System was used to maintain focus at the glass-PDMS interface. Two fields of view (FOVs) were required to image all 30 migration channels in the device.

Images were analyzed in FIJI using the following workflow: 1) Due to the high concentration of FITC-dextran (1 mg/mL) used for fluorescence detection, out-of-plane fluorescence from the chamber regions resulted in saturation along the channel edges, typically spanning 15-20 μm in length. Therefore, fluorescence intensity profiles were measured exclusively in unsaturated regions within each channel. 2) For each device, the intensity profiles of the channel closest to the source inlets, channel in the center of the device, and channel farthest from the source inlets were measured and compared over time. 3) Each intensity profile was background-subtracted and normalized to the maximum intensity within the channel region in each frame, excluding saturated areas. The gradient profile was analyzed every 1 minute up to 30 minutes after FITC-dextran introduction, then every 30 minutes for up to 5 hours.

### Cell loading into migration device

After bonding the microfluidic device to a No. 1.5 glass coverslip in a 35 mm dish (see **Design and fabrication of microfluidic migration** devices), the device was treated with high-intensity UV light for 30 minutes. A UV-sterilized syringe, needle, tubing, and stainless steel pin filled with 1X PBS were then connected to the device outlet (see **Gradient characterization in migration devices**). The device was first wetted with PBS by manual infusion of PBS into the device. PBS was then added to the device inlets and the syringe was manually withdrawn to remove any created bubbles in the device. Excess PBS was removed from the inlets and replaced with 50-60 uL of 100 μg/mL fibronectin (Sigma-Aldrich, FC010), depending on the thickness of the PDMS. The fibronectin was manually withdrawn to fill the device, needle, and some of the tubing to account for possible backflow of media back into the device during the fibronectin incubation period. The preceding method was used to exchange old media in the device, and is adopted for the rest of the procedures written in the following sections. The device was treated with fibronectin for 1.5 hours at 37 °C for the PDMS to absorb the fibronectin and become conducive for cell attachment and growth.

Cells were harvested from culture plates using 0.05% Trypsin/EDTA (Thermo Fisher Scientific Inc., 25200056) and centrifuged at 1500 rpm for 5 minutes. The cell pellet was resuspended in phenol red-free F-12 complete media at a concentration of approximately 1 × 10^5^ cells/mL. The device was washed with PBS before being filled with F-12. Excess media was removed from one of the inlets and replaced with the cell suspension. A slightly larger volume of the cell suspension was added to this inlet compared to the opposite inlet to drive cell movement into both chambers. The suspension was allowed to settle for 5 minutes before repeating this process 3-5 times until an adequate number of cells accumulated in the chamber closest to the inlet. The device was placed in a 37 °C incubator, and cell spreading was monitored every 10 minutes using a dual phase contrast/fluorescence LED microscope (Leica, PAULA).

After cell spreading, the device was taken out of the incubator and the media in the inlets were exchanged with equal volumes of serum-free phenol red-free F-12 (50-60 µL, depending on the thickness of PDMS) to induce cell cycle synchronization and heighten the cell response to growth factors (*17*). The device was then returned to the incubator and connected to a syringe pump (Harvard Apparatus, 70-3007) for 1.5 hours at a withdrawal rate of 5 µL/h. This stabilization period was imaged every 10 minutes using the Leica PAULA microscope.

### Acquisition of cell migration speeds and persistence

Following serum starvation, time-lapse imaging of cell migration under different conditions was performed using a dual phase contrast/fluorescence LED microscope (Leica, PAULA). To introduce EGF and/or endocytic inhibitors into the microfluidic device, the lid of the 35 mm dish was briefly removed and replaced after addition.

Cell migration was first imaged in the presence of an EGF gradient. After serum starvation, two pipettes were used to simultaneously introduce equal volumes of serum-free F-12 and a concentrated EGF (Thermo Fisher Scientific Inc., AF-100-15-500UG) solution into each inlet of the device (5-6 µL, depending on PDMS thickness). The EGF stock solution was prepared at a concentration that would result in a final inlet concentration of 20 ng/mL after spiking, accounting for an estimated 3.75 µL reduction per inlet volume due to prior flow. For consistency, EGF was always introduced in the inlet opposite the chamber where cells had been seeded.

Next, cell migration was imaged in the presence of an EGF gradient with 5 µM Dyngo-4a (Abcam, ab120689). After adding EGF to one inlet, two additional pipettes were used to simultaneously introduce equal volumes of serum-free F-12 and a concentrated Dyngo-4a solution into either the same inlet as EGF or the opposite inlet (<1 µL, depending on PDMS thickness). Following media addition, the dish lid was replaced, and cells were imaged every 10 minutes for up to 5 hours using the Leica PAULA system.

### Analysis of cell migration speeds and persistences

Cell migration experiments were conducted at least three times, with over 8-10 cells analyzed per condition. Time-lapse images of cell motility were analyzed over a 2-4-hour period using the FIJI plugin MTrackJ (*18*) (https://imagescience.org/meijering/software/mtrackj/). For all conditions, migration was analyzed after a 2-hour incubation period to allow Dyngo-4a treatment to take effect and to ensure flow stabilization between the two inlets. Cell trajectories were recorded by tracking the center of cell nuclei at each timepoint, and average velocities and path lengths were compiled in Excel format. Each cell’s movement was analyzed to determine when its nucleus first entered the microfluidic channel, allowing for the identification of the timepoint at which tracking commenced. Tracking was completed when the center of the nucleus exited the channel or when the time-lapse ended, whichever occurred first. Cells were selected based on the following criteria: 1) the cell was not undergoing division and 2) the cell was not obstructed by another cell in either direction of the gradient. Cell persistence was calculated as the ratio of net displacement to total path length. Differences in velocity and persistence were calculated using two-way ANOVA. A p-value of < 0.05 was considered statistically significant (see **Statistics and reproducibility**).

### Immunofluorescence imaging of migrated cells

Immediately after acquiring cell trajectories (see **Acquisition of cell migration speeds and persistence**), the entire migration device, along with its attached tubing and syringe apparatus, was carefully removed from the incubator for subsequent immunostaining. Cells were first washed three times with 1X PBS, then fixed with 3.7% paraformaldehyde (Sigma-Aldrich, P6148) for 15 minutes at room temperature on a rocker (50 rpm). After fixation, cells were washed three times with 1X PBS, and the needle, tubing, and syringe were carefully removed from the device. Permeabilization was performed using 0.1% Triton X-100 (Sigma-Aldrich, T8787) in 1X PBS for 15 minutes at room temperature, rocking. Cells were then washed three times with 1X PBS and blocked overnight at 4°C with PBS-t blocking buffer (1X PBS containing 2% (v/v) rat serum (Thermo Fisher Scientific Inc., 10-710-C) and 0.1% Tween-20 (Sigma-Aldrich, P9416)), while gently rocking (30 rpm). The following day, cells were incubated for 1 hour at room temperature with paxillin primary antibody (Thermo Fisher Scientific Inc., AHO0492) diluted 1:500 in 1% (v/v) rat serum and 0.1% Tween-20 in PBS. After three PBS-t washes, cells were incubated for 45 minutes at room temperature, protected from light, with fluorochrome-conjugated secondary antibody (Thermo Fisher Scientific Inc., A-21124) diluted 1:3000 in 1% (v/v) goat serum (SouthernBiotech, 0060-01) and 0.1% Tween-20 in PBS. Finally, nuclei were stained using Hoechst dye (1:2000 dilution in PBS-t, Thermo Fisher Scientific Inc., H21486) for 10 minutes at room temperature, rocking. Cells were washed three times with PBS-t and stored in 1X PBS for imaging.

Cells were imaged using a spinning disk confocal microscope (see **Spinning disk confocal image acquisition**), equipped with a 100X objective lens (NA 1.45, Nikon) with 405 nm excitation (300 ms exposure time, 50% power), 488 nm excitation (300 ms exposure time, 50% power), and 561 nm excitation (300 ms exposure time, 100% power). To capture AP-2 and paxillin throughout the entire cell volume, images were acquired across 11-13 planes at 0.5 μm intervals. The Nikon Perfect Focus System was used to maintain focus at the interface between the glass and PDMS. Additionally, images were collected 1 μm below the perfect focus position to ensure optimal imaging of the cell’s basal plane.

Cells were selected for imaging based on the following criteria: 1) the entire cell body remained within a single migration channel, 2) the cell was not undergoing division, and 2) the cell was not obstructed by another cell in either direction of the gradient.

### Spinning disk confocal image acquisition

Imaging was performed using a Nikon Ti-E fluorescence microscope equipped with a CSU-W1 spinning disk unit (Yokogawa Electric Corporation), a 100X Plan-Apochromat Lambda objective (1.45 NA, Nikon), an sCMOS camera (Prime 95B; Teledyne Photometrics), and 405 nm, 488 nm, and 561 nm excitation lasers. Three-dimensional images of AP-2, paxillin, and cell nuclei were acquired using NIS Elements software.

### Analysis of focal adhesions and clathrin pits

Following imaging (see **Immunofluorescence imaging of migrated cells**), confocal z-stacks were processed for quantitative analysis of paxillin and AP-2-labeled endocytic structures in FIJI. The paxillin and AP-2 channels underwent background subtraction, after which images were converted to 8-bit format for further processing. Maximum intensity projections (MIP) were generated for both channels.

To define the front and rear regions of migrating cells, the balloon segmentation plugin (https://imagej.net/plugins/balloon) was applied to identify the cell boundary based on intensity gradients. A custom FIJI macro then iteratively adjusted the segmentation to ensure equal area partitioning between the front and rear. The script saved the segmented regions as regions of interest (ROIs), which were then used to quantify front-to-rear differences in protein distributions.

Moments thresholding was applied to both channels to segment fluorescence signals. The AP-2 channel underwent additional watershed analysis to separate overlapping structures, while the paxillin channel was analyzed without further segmentation due to its diffuse appearance. The particle sizes of AP-2 puncta were measured within the front and rear regions. Paxillin and AP-2 intensities were assessed by normalizing the integrated density of the front and rear regions to the whole-cell intensity.

### Total internal reflection fluorescence structured illumination microscope (TIRF-SIM) image acquisition

TIRF-SIM images were acquired using a custom-built system previously described (*19*). This system is constructed on an inverted Nikon Eclipse TI-E microscope body, equipped with an Olympus APO 100× 1.49 NA objective and a Hamamatsu ORCA-Fusion BT sCMOS camera. Structured laser illumination (488 nm or 561 nm, 300 mW; Coherent, SAPPHIRE LP) is produced via an acousto-optic tunable filter (AOTF; AA Quanta Tech, AOTFnC-400.650-TN), a polarizing beam splitter, an achromatic half-wave plate (HWP; Bolder Vision Optik, BVO AHWP3), and a ferroelectric spatial light modulator (SLM; Forth Dimension Displays, QXGA-3DM-STR). Images acquired with varying phases and orientations are subsequently reconstructed into a super-resolution image using the HiFi-SIM algorithm (*20*).

### Statistics and reproducibility

All data presented reflect at least three biologically independent experiments per experimental condition. Statistical analyses and graphical representations were performed using RStudio and GraphPad Prism.

Graphical data representations and statistical analyses are described in the respective figure legends. Statistical significance was assessed using two-way ANOVA, Dunn’s multiple comparisons test, or two-sided Mann-Whitney U test, as appropriate. P-values are reported as follows: p > 0.05, *: p ≤ 0.05, **: p ≤ 0.01, ***: p ≤ 0.001, ****: p ≤ 0.0001.

## Results and discussion

### A stable EGF gradient exists across all microchannels

We designed and fabricated our platform to generate spatially linear biochemical gradients across confined, parallel microchannels while minimizing flow-mediated mechanical stimulation (**Fig. 1A**) (*21*). At a low flow rate of 5 µL/h, COMSOL simulations confirmed that fluid flow remained restricted to the chambers of the device, with negligible velocity in the migration channels (**Fig. S1A**). A z-profile of velocity magnitude across all 30 channels demonstrated that flow at the channel midplane was nearly zero (**Fig. S1B and C**). We estimated shear stress in the chambers to be approximately 0.05 dyn/cm^2^, which is well below the threshold reported to induce migration through flow-dependent mechanisms, indicating that cell migration in this system is governed by the imposed biochemical gradient rather than mechanical cues (*22*).

**Fig. 1.**
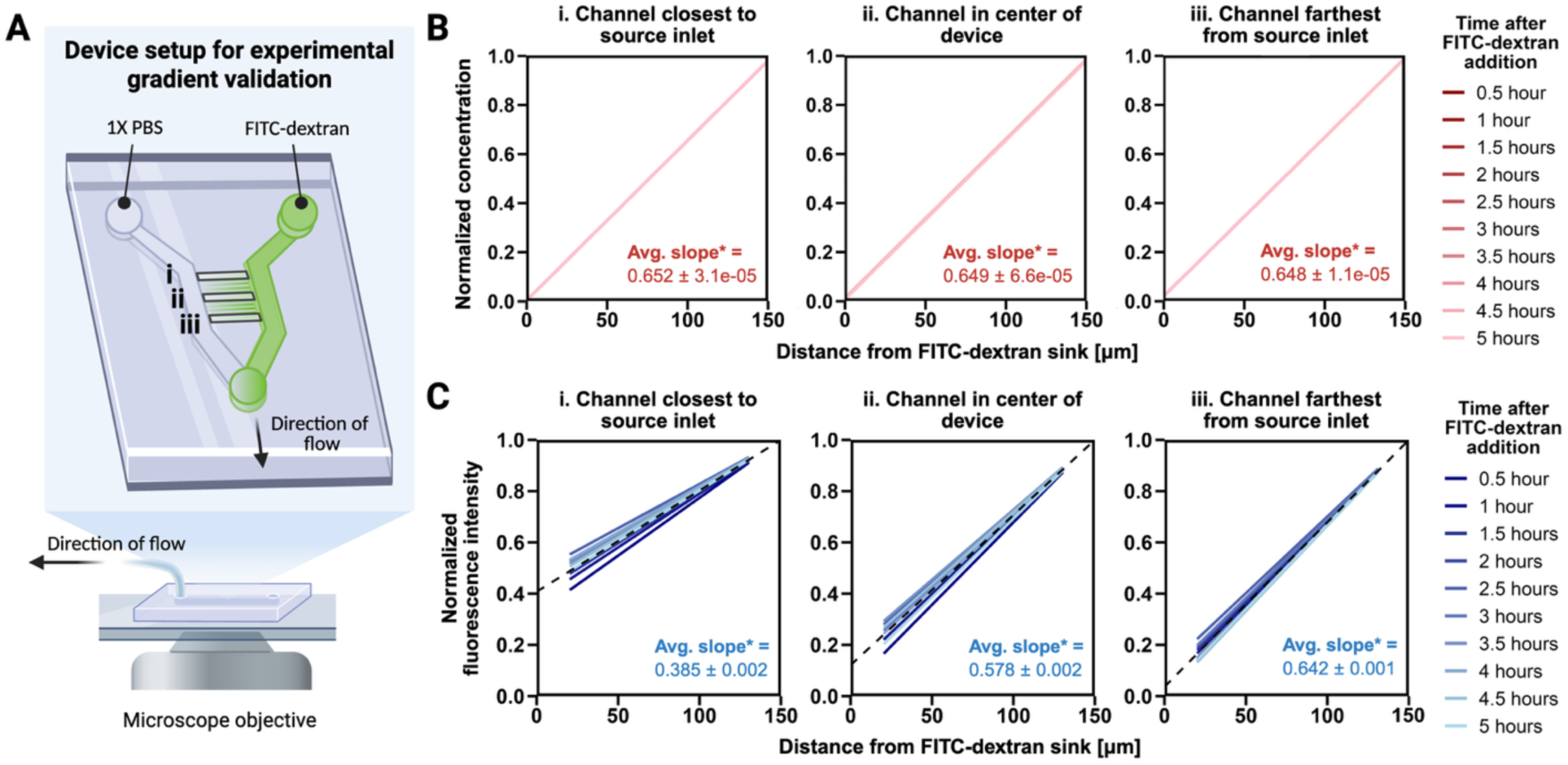
Validation of stable FITC-dextran gradients across microfluidic migration channels. (A) Experimental setup illustrating 10 kDa FITC-dextran introduced through the source inlet and sampled at defined channels across the device (i: closest to the source inlet, ii: center, iii: farthest from the source inlet). Flow is maintained using a syringe pump at a low flow rate, and the microscope objective is positioned under the migration channels. (B) Simulated gradients showing representative normalized concentration profiles at three channel positions, plotted against distance from the FITC-dextran sink. *Slopes are scaled by a factor of 100 for readability. (C) Experimental gradients showing normalized fluorescence intensity of FITC-dextran plotted against distance from the sink at three channel positions (N=3). Data are truncated at the channel entrance and exit due to high out-of-plane fluorescence. The dashed black line represents the average linear fit across all timepoints. *Slopes are scaled by a factor of 100 for readability.

To characterize the physical structure of the platform, we used scanning electron microscopy to confirm key microdevice dimensions, including a channel height of ∼5 µm, width of 20 µm, and the spacing profile between adjacent channels (**Fig. S2A**). We then performed optical calibration using 10 kDa FITC-dextran, selected for its comparable molecular weight to epidermal growth factor (EGF, ∼6 kDa), and observed a linear relationship between fluorescence intensity and concentration (**Fig. S2B**), enabling precise quantification of gradient strength using fluorescence imaging. COMSOL simulations predicted that a steady-state gradient would form within 3 minutes, with consistent and steep linear concentration profiles across representative channels (**Fig. 1B, Fig. S1D**, **Movie S1**). Time-lapse imaging confirmed that experimental gradients formed within 5 minutes and remained stable by 30 minutes, although the resulting slopes were shallower than those predicted by simulation (**Fig. 1C, Fig. S2C**). The gradient was most shallow in the channel closest to the source inlet and gradually steepened across the device, closely matching the simulated slope in the channel furthest from the source. These differences likely arise from experimental factors not captured in the simulation, such as minor asymmetries in device geometry and mixing delays introduced by manual addition of FITC-dextran. Despite this, the device reliably produced spatially stable gradients across all channels. For downstream migration analyses, we focused on cells in the middle to bottom 15 channels of the microdevice, where experimental gradients most closely aligned with model predictions.

### Microdevice design supports high-resolution imaging of subcellular structures during directed cell migration

A key attribute of our migration platform is its compatibility with high-resolution and super-resolution microscopy, enabling direct visualization of subcellular structures during chemotactic migration. We engineered the microchannel heights to be 5 μm, which both constrains cell morphology for axial alignment and permits optical access from below the device using high numerical aperture objectives (**Fig. 2A and B**). This allowed us to confirm that cells migrating through the channels remain within the working distance of oil-immersion lenses used in high resolution spinning disk confocal and total internal reflection fluorescence structured illumination microscopy (TIRF-SIM).

**Fig. 2.**
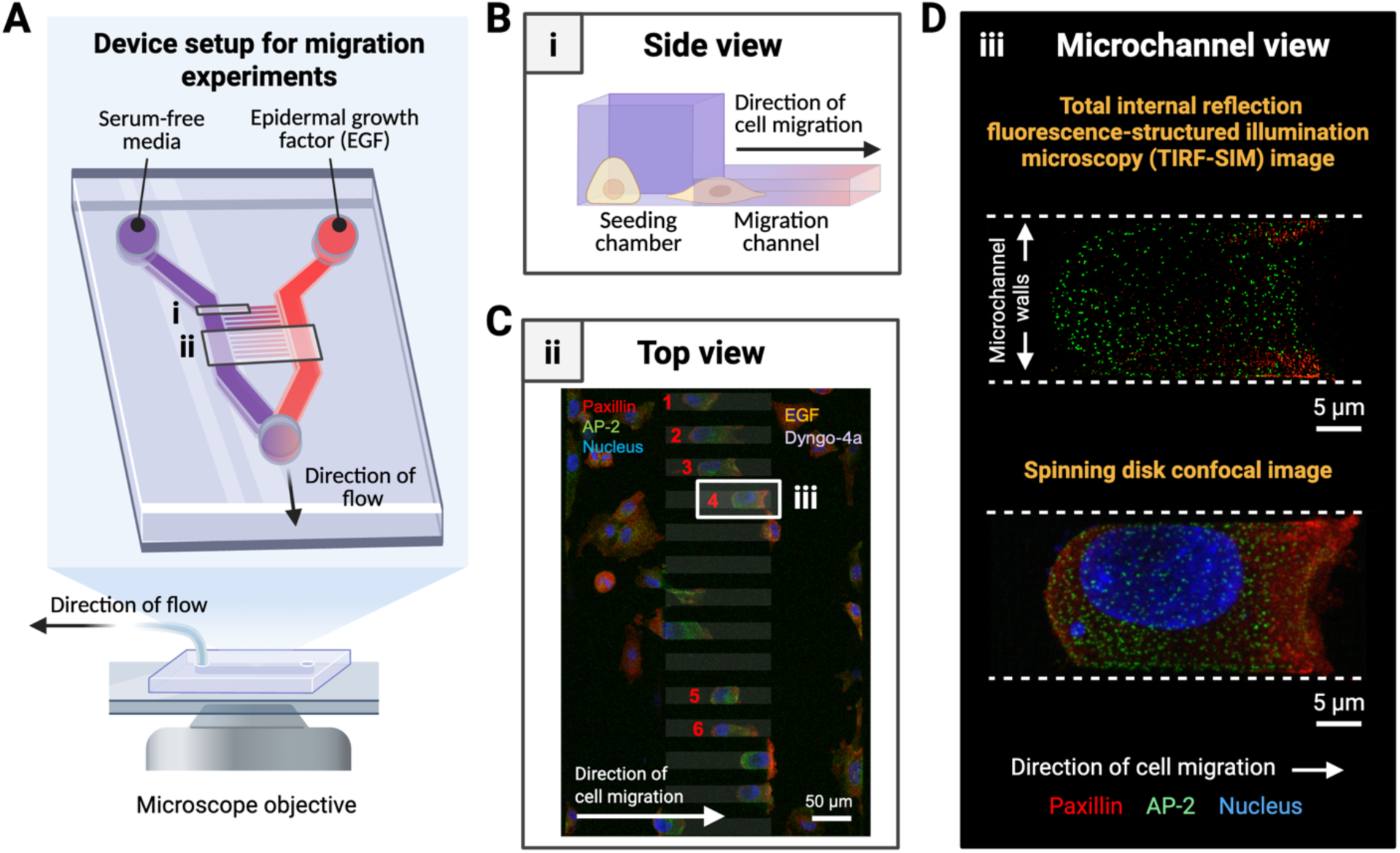
Device design and imaging modalities for characterizing cancer cells during migration. (A) Experimental setup of the microfluidic device used for migration assays, showing the introduction of EGF through the source inlet and the microscope objective positioned beneath the migration channels. Two representative regions are indicated by (i) and (ii), corresponding to panels (B) and (C), respectively. (B) Side view of the microchannel structure, highlighting the height difference between the seeding chamber and the 5 μm-tall migration channels. (C) Maximum intensity projection showing a top view of a portion of the device in an experiment where Dyngo-4a was applied to the same chamber as EGF. Cells endogenously express AP2-eGFP (green) and are immunostained for paxillin-mCherry (red) and Hoechst for the nucleus (blue). Tracked cells (6) are indicated by red numbers. Arrow indicates the direction of cell migration. A representative region indicated by (iii) corresponds to panel (D). Scale bar: 50 μm. (D) Maximum intensity projections of representative migrating cells acquired using TIRF-SIM (top) and spinning disk confocal microscopy (bottom). Cells are stained for paxillin (red) and the nucleus (blue), and endogenously express AP2-eGFP (green). Dashed lines indicate the boundaries of the migration channel walls. Arrow indicates direction of cell migration. Scale bar: 5 μm.

We tested this compatibility by imaging migrating cells expressing endogenous AP2-eGFP (a marker for CME structures), immunostained for paxillin-mCherry (a focal adhesion marker), and dyed with nuclear Hoechst (**Fig. 2C, Fig. S3A**). Both TIRF-SIM and spinning disk confocal modalities produced high-resolution images within the channel confines, resolving distinct spatial distributions of adhesion and endocytic proteins (**Fig. 2D, Fig. S3B and C, Movie S2**). In cells located outside the channels, AP-2 and paxillin displayed more uniform distributions, and polarization was less apparent (**Fig. S3B and C**). These observations underscore how microchannel confinement and chemotactic cues enhance spatial polarization of adhesion and endocytic machinery. Since we aim to visualize adhesive and endocytic structures throughout the full depth of the cell, for the remainder of the text, we focus on cells imaged in 3D using spinning disk confocal microscopy.

### Front-rear endocytic inhibition reveals spatial control of paxillin and AP-2 distribution

To examine how endocytic activity influences adhesion dynamics during directed migration, we applied Dyngo-4a, a small molecule inhibitor of dynamin-mediated CME (*23–25*), to either the front or rear chamber of the microfluidic device. This spatially selective delivery approach allowed us to perturb specific regions of the cell without altering the entire pericellular microenvironment. The ability to restrict drug exposure to the leading or trailing edge is primarily enabled by the physical confinement of cells within the channels, which limits molecular diffusion around the cell body.

Following gradient exposure and migration (**Movie S3**), we fixed and immunostained endogenously expressing AP2-eGFP cells for paxillin and nuclei (**Fig. 3A**), then analyzed them using a custom front-rear segmentation pipeline (**Fig. S4**). High-resolution imaging revealed that inhibiting endocytosis at either end of the cell disrupted the polarized distribution of paxillin and AP-2 fluorescence (**Fig. 3B and C**). Quantification showed that paxillin remained front-enriched under both control conditions and front-targeted Dyngo-4a treatment (**Fig. 3B**). In contrast, rear-targeted inhibition eliminated this front-rear asymmetry, resulting in no significant difference in paxillin intensity between the leading and trailing edges. Similarly, AP-2 was front-enriched under control and front-targeted Dyngo-4a treatment, but rear-targeted inhibition led to a uniform distribution of both proteins (**Fig. 3C**).

**Fig. 3.**
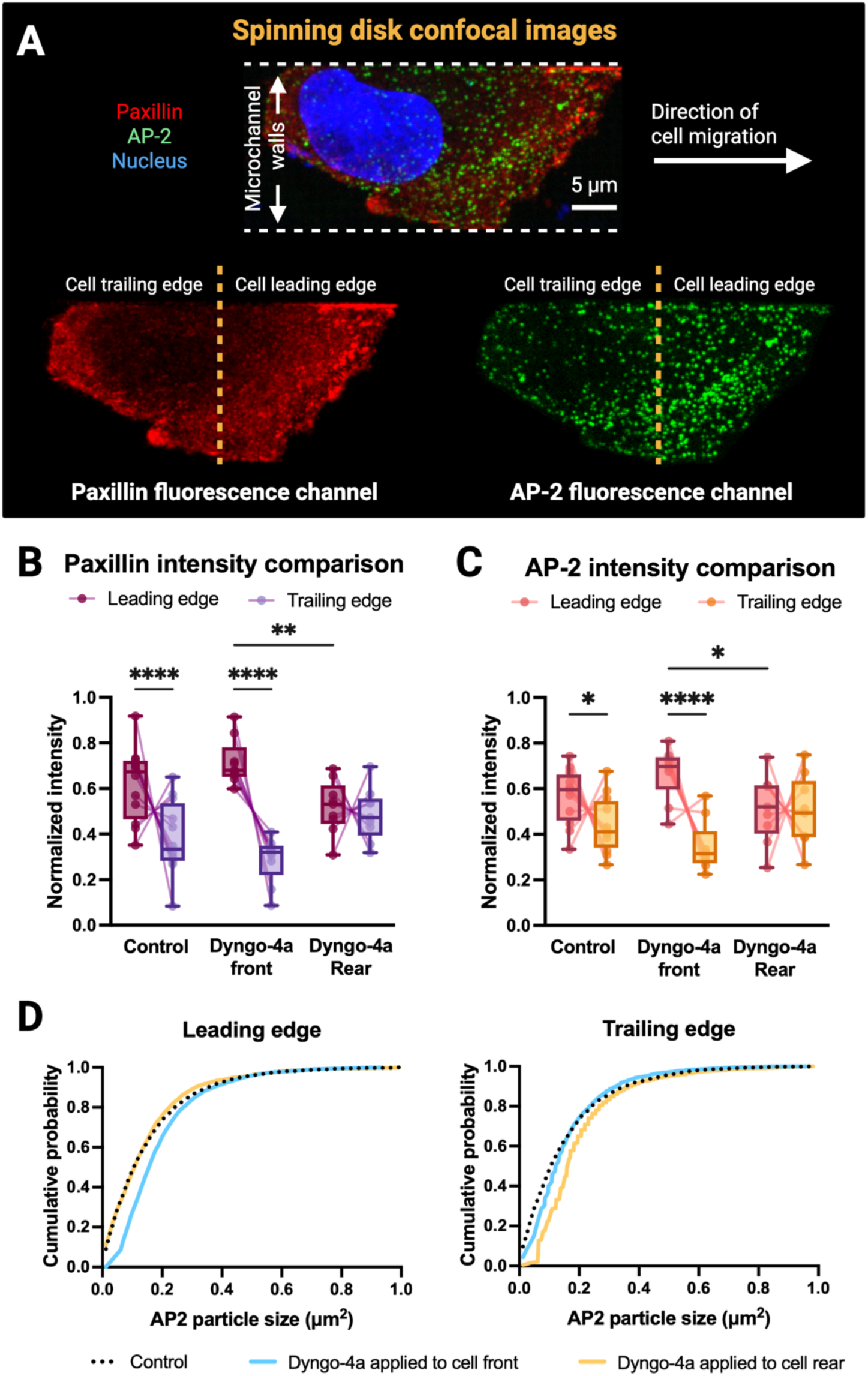
Inhibition of endocytosis alters the front-rear distribution of paxillin and AP-2 structures, as well as the size of AP-2 particles. (A) Maximum intensity projection of spinning disk confocal images showing a migrating cell within a microchannel, displaying paxillin (red), AP-2 (green), and the nucleus (blue). White dashed lines indicate the microchannel walls; yellow dashed lines denote the division between the cell leading and trailing edges. Arrow indicates direction of cell migration. Scale bar: 5 μm. (B) Quantification of normalized paxillin fluorescence intensity at the cell leading and trailing edges under control conditions and with Dyngo-4a applied to the front or rear of the cell. (C) Quantification of normalized AP-2 fluorescence intensity under the same conditions as (B). (D) Cumulative distribution plots of AP-2 particle sizes at the leading and trailing edges. All pairwise comparisons are significant (P < 0.001), except for Control vs. Dyngo-4a applied to the rear at the leading edge, and Control vs. Dyngo-4a applied to the front at the trailing edge. Box plots indicate the median, 25th, and 75th percentiles; each dot represents a single cell, with lines connecting leading and trailing edge values. Sample sizes: Control (N = 3, n = 12), Dyngo-4a applied to front (N = 3, n = 9), Dyngo-4a applied to rear (N = 3, n = 9). *P < 0.05, **P < 0.01, ***P < 0.001, ****P < 0.0001.

Particle-level analysis of AP-2 puncta showed that Dyngo-4a treatment also altered particle size distributions (**Fig. 3D**). Cells exposed to Dyngo-4a at the front exhibited a rightward shift in particle size at the leading edge, consistent with an accumulation of larger endocytic structures. A similar effect was observed when Dyngo-4a was applied to the rear. This pattern likely reflects the mechanism of Dyngo-4a, which inhibits scission of clathrin-coated pits but not their formation or growth (*24*, *26*), resulting in stalled and enlarged endocytic intermediates. These localized accumulations indicate that endocytic activity is spatially regulated and suggest that active internalization is required to maintain the polarized distribution and turnover of adhesion and endocytic complexes during migration.

### Endocytic inhibition differentially impacts migration speed and persistence depending on the site of application

We lastly investigated how spatially controlled inhibition of endocytosis affects migration dynamics by quantifying speed and persistence in migrating cells under control conditions and with Dyngo-4a applied to either the front or rear of the cell. These measurements rely on cells not blocking the microchannels at either end, ensuring that the inhibitor exposure remains spatially restricted. Front-targeted Dyngo-4a treatment significantly increased both migration speed and persistence compared to control, whereas rear-specific treatment reduced speed without affecting persistence (**Fig. 4A and B**). These findings suggest that endocytosis at the leading edge of single cells may act as a regulatory brake on cell motility, while endocytosis at the trailing edge supports sustained motility.

**Fig. 4.**
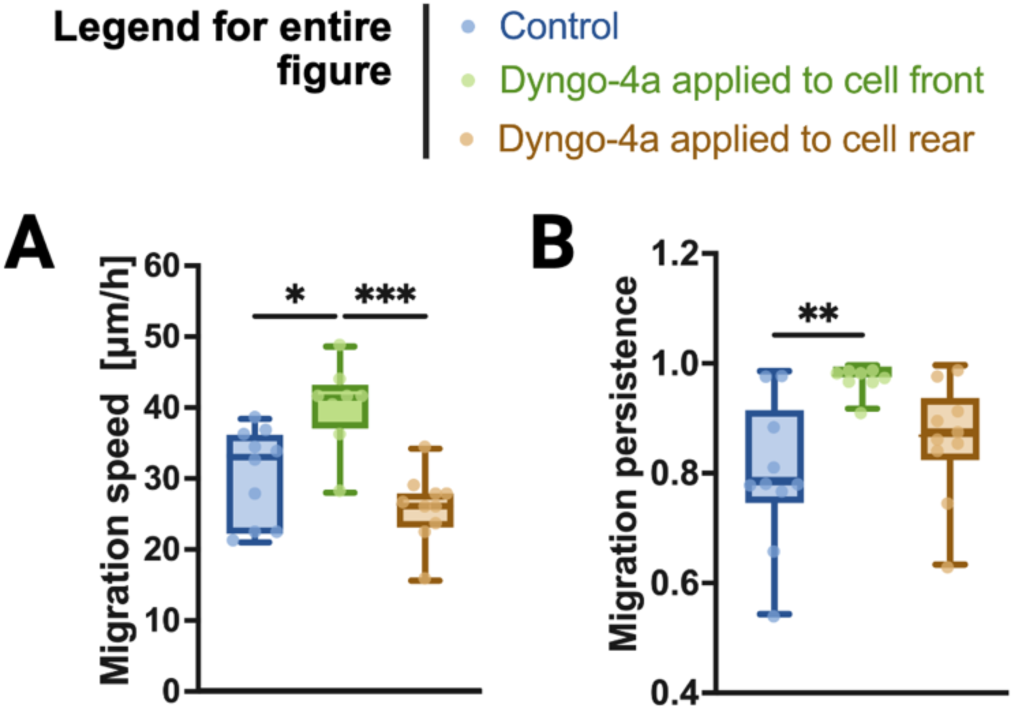
Inhibition of endocytosis alters migration speed and persistence in single cells. (A) Migration speed of cells under control conditions, or with Dyngo-4a applied to the cell front or rear. (B) Migration persistence of cells under the same conditions as in (A). Persistence is defined as the ratio of net displacement to total path length, with values closer to 1 indicating more directed migration. Box plots show median, 25th, and 75th percentiles; each dot represents an individual cell trajectory. Statistical significance: *P < 0.05, **P < 0.01, ***P < 0.001, ****P < 0.0001. Sample sizes (all N = 3): Control (n = 10), Dyngo-4a applied to cell front (n = 8), and Dyngo-4a applied to cell rear (n = 10).

## Conclusion

Targeting the spatial regulation of endocytosis offers a new strategy to understand how cells navigate confined environments, particularly in the context of cancer invasion. Building on previous studies that implicate the role of endocytosis in adhesion turnover and migratory plasticity (*27–29*), we hypothesized that locally inhibiting CME would disrupt the polarized distribution of adhesion and endocytic machinery. Our findings support the hypothesis that front-targeted inhibition of endocytosis led to increased accumulation of paxillin and AP-2 at the leading edge, while rear-targeted inhibition reduced their front-enriched localization. Notably, rear-targeted inhibition was more effective in eliminating polarity. This result suggests that endocytosis at the cell rear may be especially critical for recycling membrane components and clearing trailing-edge adhesions, which are processes that reinforce directional front-rear asymmetry necessary to maintain migration in mechanically constrained spaces (*28*, *30–32*).

In line with this outcome, we observed that front-targeted endocytic inhibition increased migration speed and directional persistence. Rear-targeted inhibition reduced speed but did not significantly affect persistence. These spatially distinct outcomes suggest that CME plays different roles at the front and rear of migrating cells. One possible explanation is that CME facilitates the removal of activated receptors or adhesion complexes, which may otherwise act as traction points or molecular brakes. Disruption of this process at the cell front could lead to more unrestrained cell movement, enhancing both speed and directional persistence.

Together, these results underscore the utility of our microfluidic system in dissecting how localized molecular events govern cell migration in a physiologically relevant environment. By enabling targeted drug delivery to subcellular regions of migrating cells, our platform offers a valuable tool for probing spatially resolved signaling processes. This work highlights that CME plays distinct roles in modulating speed versus persistence, depending on both cellular context and spatial directionality. Live-cell imaging of AP-2 and paxillin during migration in this system will be especially informative, as prior work has shown that focal adhesions and clathrin-containing adhesion complexes (a distinct class of adhesive structures associated with clathrin and endocytic machinery) coexist but follow different temporal patterns in SUM-159 breast cancer cells, influencing both cell cycle progression and potentially migratory behavior (*33*). Future studies will be needed to determine the downstream signaling consequences of polarized endocytosis and whether specific cargo internalization events are responsible for the observed migratory phenotypes. More broadly, our findings suggest that spatial control of trafficking may be a viable therapeutic target to disrupt single-cell tumor escape without necessarily impairing normal tissue maintenance or immune surveillance.

## Data availability

All data and custom scripts used in this study are available upon reasonable request.

## Author Contributions

E.T.C. designed, validated, and fabricated the confined migration devices, conducted migration experiments, performed immunostaining, acquired spinning disk confocal microscopy images of the stained samples, and performed data analysis. E.T.C. and T.H.J. optimized the experiments and performed COMSOL modeling of gradient establishment in the migration device. C.M.T. imaged immunostained cells using TIRF-SIM and performed image reconstruction. H.K. and M.W.D. measured cell migration speeds and persistences in microdevices and analyzed immunofluorescence images. M.W.D. and M.M.A. acquired spinning disk confocal microscopy images of cells. E.T.C., J.W.S., and C.K. oversaw the project and prepared the manuscript. All authors discussed the results and contributed to revisions of the manuscript.

## Acknowledgements

The authors thank all past and present members of Jonathan Song’s and Comert Kural’s labs for critical discussions about experimental design. E.T.C. would like to thank Keith Ramsey, Kavya Dathathreya, and Dave Hollingshead from Ohio State’s Nanotech West laboratory, in addition to Melikhan Tanyeri from Duquesne University for their assistance in developing and characterizing the migration devices. E.T.C. was supported by the Ohio State University Molecular Biophysics Training Program (NIH T32 GM118291-04) and the Pelotonia Fellowship Program. H.K. was supported by The Ohio State University Beckman Scholars Program and Department of Chemistry and Biochemistry. M.W.D. was supported by the Stamps Eminence Scholarship Program at The Ohio State University. M.M.A. was supported by the Ronald E. McNair Postbaccalaureate Achievement Program at The University of Wisconsin, Madison. C.K. was supported by NSF Faculty Early Career Development Program (award number: 1751113) and NIH R01GM127526. J.W.S. was supported by NSF CBET-1752106. Any opinions, findings, and conclusions expressed in this material are those of the authors and do not necessarily reflect those of the Pelotonia Fellowship Program or The Ohio State University. All figures were made with BioRender.com.

