## Supplemental figures for "A microfluidic gradient and parallel-track system uncovers spatial control of endocytosis and adhesion formation in breast cancer cell migration"

### Fig. S1

COMSOL modeling and validation of gradient formation in the microfluidic device.

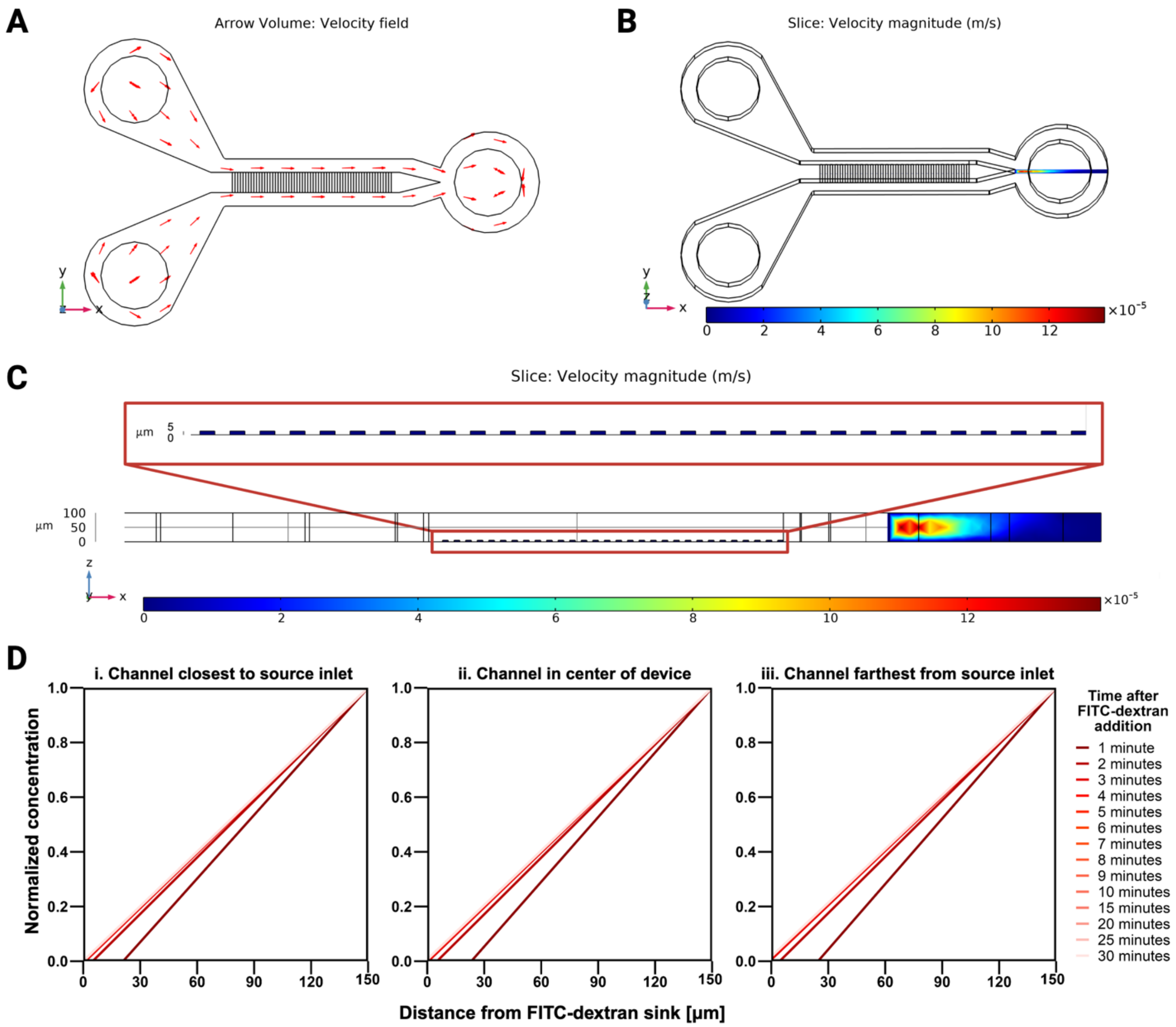


(A) Simulated velocity field showing flow streamlines through the inlet, chambers, and outlet, with no flow in the microchannels.

(B) Location of the midplane slice used for the velocity magnitude simulation shown in (C).

(C) Z-profile slice through the migration channel array showing velocity magnitude across all 30 channels. Velocity approaches zero at the channel midplane, indicating a stable no-flow environment.

(D) Simulated validation of FITC-dextran gradient formation over time at three representative channel positions: (i) closest to the source inlet, (ii) center of the device, and (iii) farthest from the source inlet. Normalized concentration is plotted against distance from the FITC-dextran sink, showing progressive gradient development over 30 minutes post-injection.

**Fig. S2**

Structural characterization and optical calibration for quantitative gradient analysis in microfluidic assays.

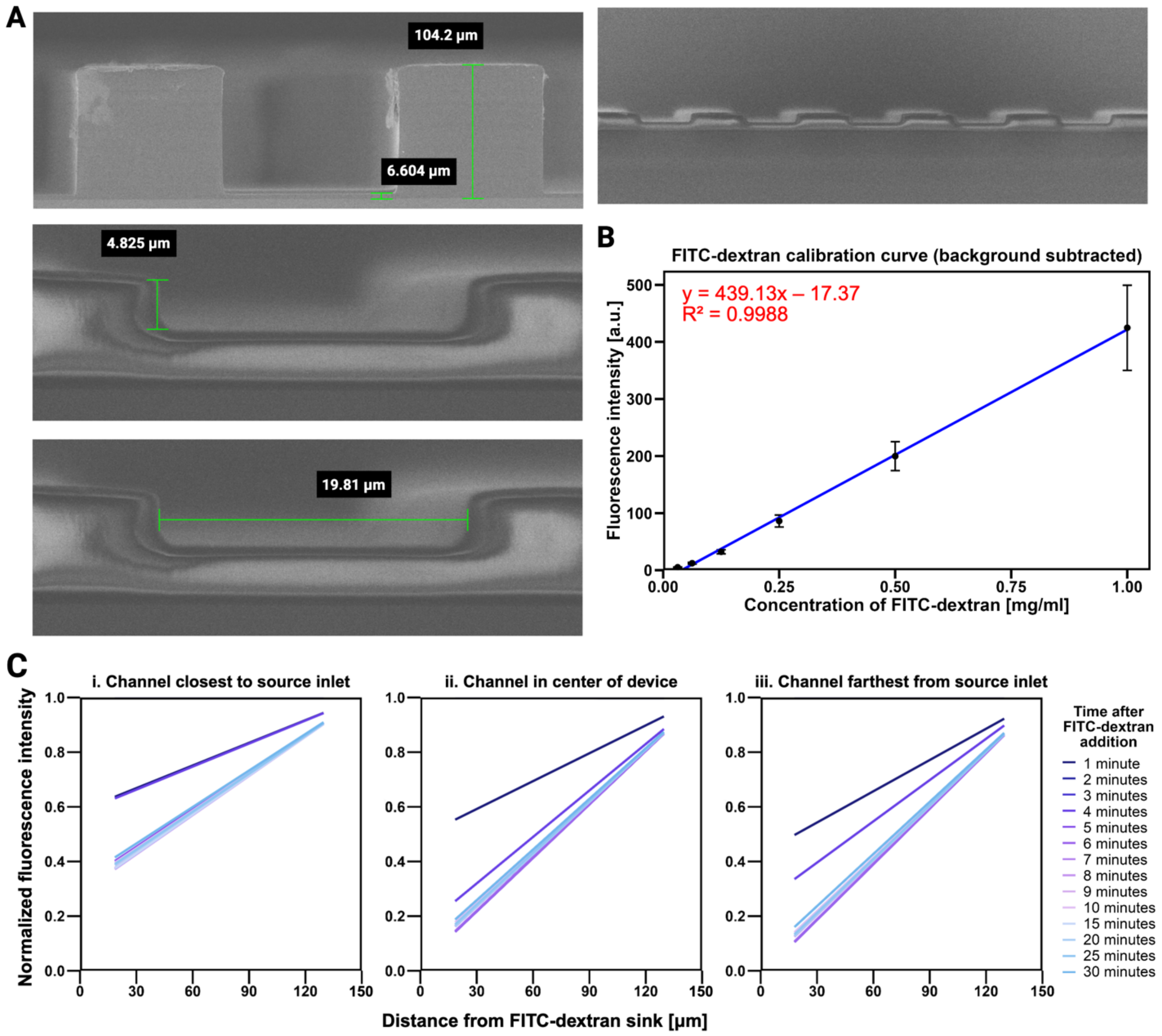


(A) Scanning electron microscopy images of the microfluidic device showing key dimensions: chamber height (top left), channel height (middle left), channel width (bottom left), and an overview of the multiple-channel array (right). The middle and bottom left images, as well as the right image, were taken from wafer regions that were broken after metal coating, resulting in visible reflectance from the SU-8 sidewalls.

(B) Calibration curve of FITC-dextran fluorescence intensity (background subtracted) as a function of concentration. Data are fit with a linear regression with $R^2$ overlaid on the graph.

(C) 30-minute time-course of FITC-dextran gradient formation normalized across three representative channels: (i) closest to the source inlet, (ii) center of the device, and (iii) farthest from the source inlet (N = 3).

**Fig. S3**

TIRF-SIM imaging of cells outside of migration channels.

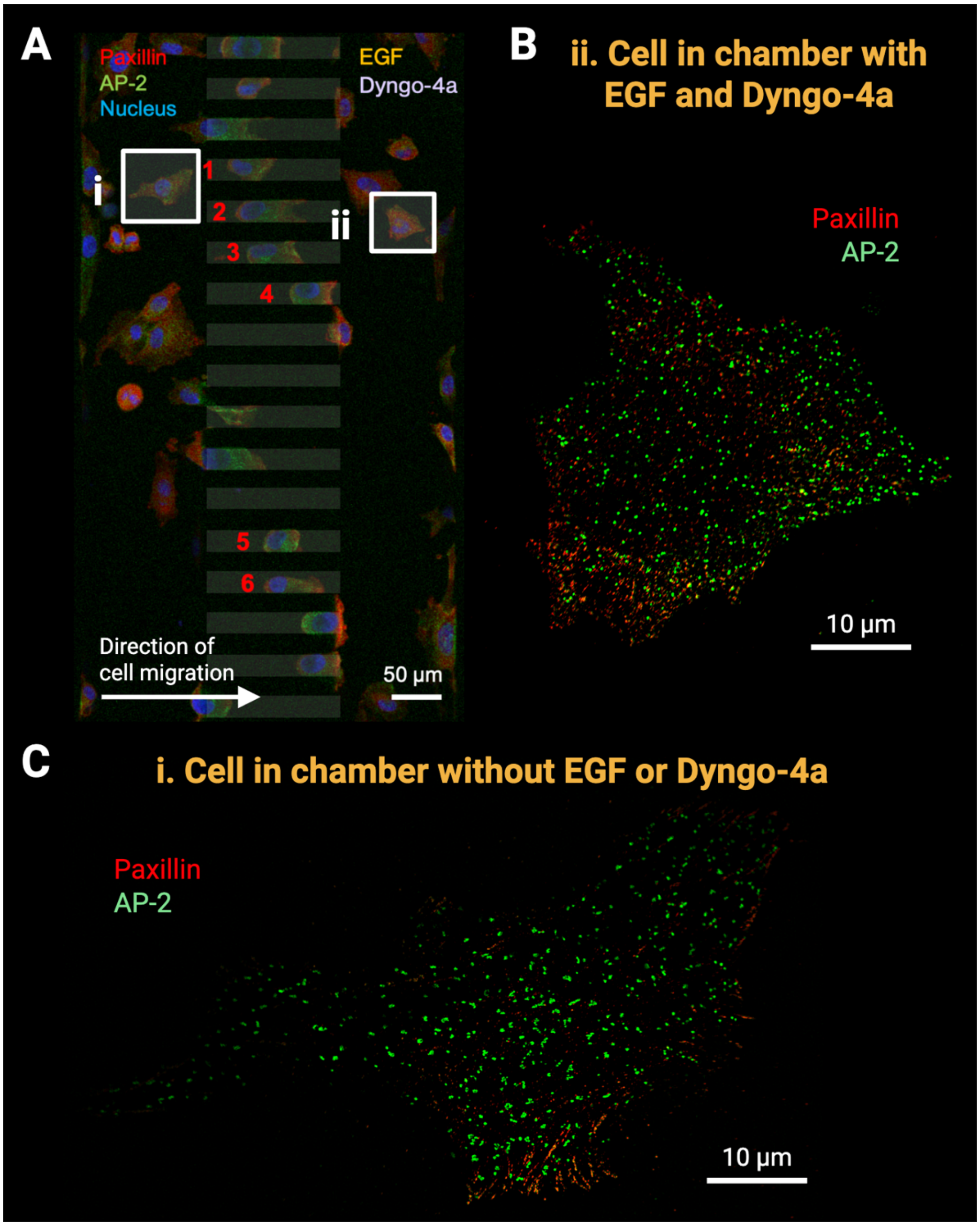


(A) Maximum intensity projection showing a top view of a portion of the device in an experiment where Dyngo-4a was applied to the same chamber as EGF (same image as shown in **Fig. 2C**). Cells endogenously express AP2-eGFP (green) and are immunostained for paxillin-mCherry (red) and Hoechst for the nucleus (blue). Tracked cells (6) are indicated by red numbers. Arrow indicates the direction of cell migration. Two representative regions are shown: (i) corresponds to panel (B); (ii) corresponds to panel (C). Scale bar: 50 µm.

(B) Cell in chamber with EGF and Dyngo-4a added. AP-2 and paxillin are broadly distributed across the cell. Scale bar: 10 µm.

(C) Cell in chamber with no EGF or Dyngo-4a added. AP-2 localizes to puncta concentrated in the midsection, and filopodia are visible at the periphery. Scale bar: 10 µm.

**Fig. S4**

Stepwise image analysis workflow for quantifying front-rear localization of paxillin and AP-2.

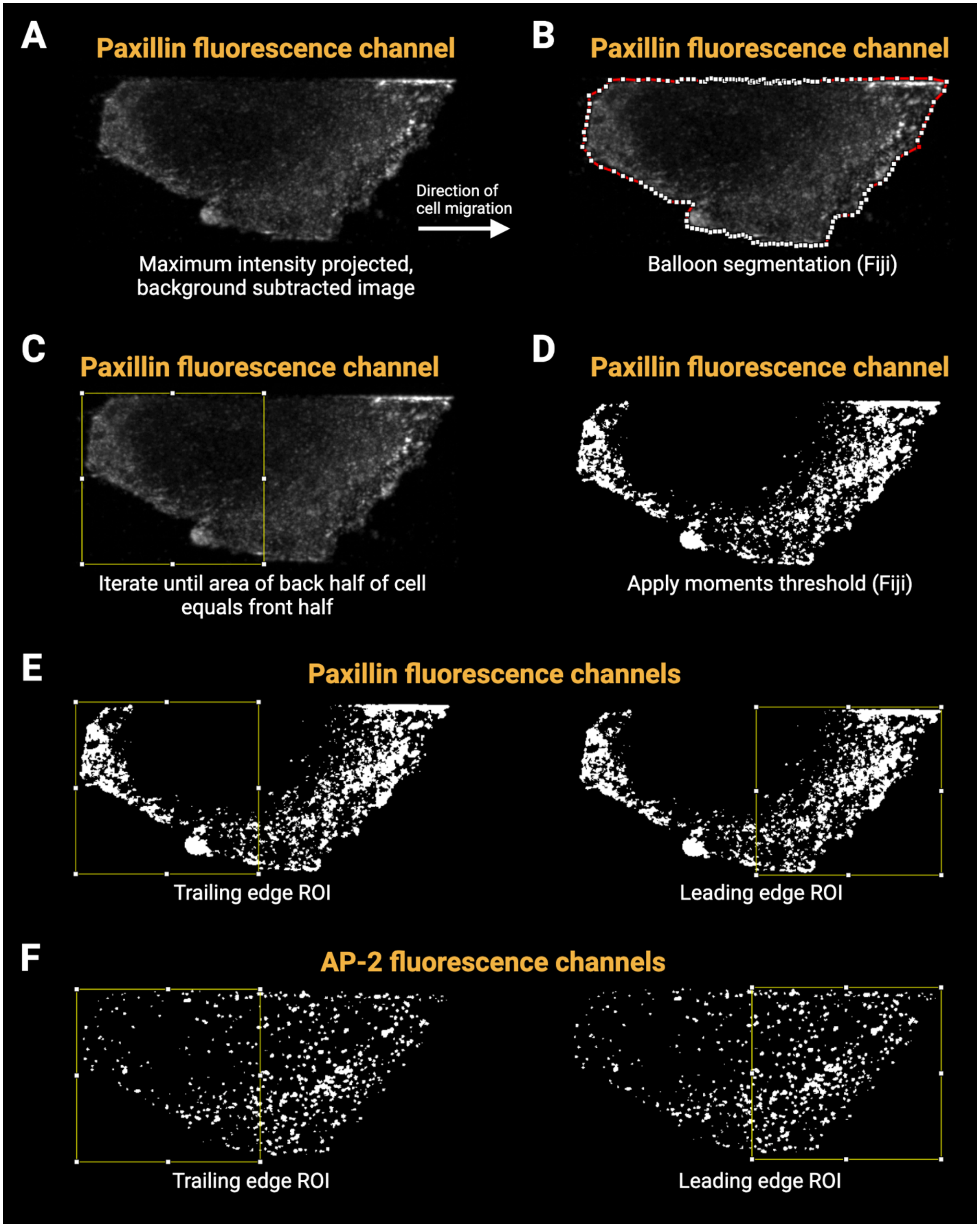

(A) A maximum intensity projection of the background-subtracted paxillin fluorescence channel is generated in FIJI.

(B) Balloon segmentation is applied to define the cell boundary.

(C) The cell is divided into front and rear halves by iteratively adjusting the division until both regions have equal area using an automated script.

(D) The moments threshold is applied to isolate mature paxillin structures.

(E) Segmented paxillin signal displayed within the trailing and leading edge-ROIs.

(F) The same ROIs are applied to the AP-2 fluorescence channel for downstream front-rear quantification, which includes watershed analysis to separate overlapping AP-2 structures.
