## Supplemental movies captions for "A microfluidic gradient and parallel-track system uncovers spatial control of endocytosis and adhesion formation in breast cancer cell migration"

### Supplemental movie captions

#### Movie S1

COMSOL simulations were performed using a molecule with the same parameters as 10 kDa FITC-dextran (diffusion coefficient = $1.43 \times 10^{-10}$ $m^2/s$). The molecule was introduced into the top chamber of the device at a concentration of 0.1 $mol/m^3$, with flow maintained through the outlet at 5 µL/h.

#### Movie S2

This movie shows the same cell featured in **Fig. 3A** and **Fig. S4A-F**, imaged across 12 z-planes at 0.5 µm intervals to capture the full cell volume within a microchannel (z-positions labeled in the top left). The 0 µm plane corresponds to the ventral membrane-glass interface, and the 4.5 µm plane represents the dorsal membrane in contact with the fibronectin-coated PDMS ceiling. The cell endogenously expresses AP2-eGFP (green) and is immunostained for paxillin-mCherry (red) and Hoechst for the nucleus (blue). Scale bar: 5 µm.

#### Movie S3

The full sequence of cell seeding, serum starvation, migration, and cell tracking in the microdevice, corresponding to the same experiment as displayed in **Fig. 2C-D**, **Fig. S3A**. The timestamp is displayed in minutes. In this experiment, EGF and Dyngo-4a were added to the right chamber. Scale bar: 100 µm.
